# Insights from the revised complete genome sequences of *Acinetobacter baumannii* strains AB307-0294 and ACICU belonging to global clone 1 and 2

**DOI:** 10.1101/641779

**Authors:** Mohammad Hamidian, Ryan Wick, Rebecca M. Hartstein, Louise Judd, Kathryn E. Holt, Ruth M. Hall

## Abstract

2.

The *Acinetobacter baumannii* global clone 1 (GC1) isolate AB307-0294, recovered in the USA in 1994, and the global clone 2 (GC2) isolate ACICU, isolated in 2005 in Italy, were among the first *A. baumannii* isolates to be completely sequenced. AB307-0294 is susceptible to most antibiotics and has been used in many genetic studies and ACICU belongs to a rare GC2 lineage. The complete genome sequences, originally determined using 454 pyrosequencing technology which is known to generate sequencing errors, were re-determined using Illumina MiSeq and MinION (ONT) technologies and a hybrid assembly generated using Unicycler. Comparison of the resulting new high-quality genomes to the earlier 454-sequenced version identified a large number of nucleotide differences affecting protein coding features, and allowed the sequence of the long and highly-repetitive *bap* and *blp1* genes to be properly resolved for the first time in ACICU. Comparisons of the annotations of the original and revised genomes revealed a large number of differences in the protein coding features (CDSs), underlining the impact of sequence errors on protein sequence predictions and core gene determination. On average, 400 predicted CDSs were longer or shorter in the revised genomes and about 200 CDS features were no longer present.

**Impact statement:** The genomes of the first 10 *A. baumannii* strains to be completely sequenced underpin a large amount of published genetic and genomic analysis. However, most of their genome sequences contain substantial numbers of errors as they were sequenced using 454 pyrosequencing, which is known to generate errors particularly in homopolymer regions; and employed manual PCR and capillary sequencing steps to bridge contig gaps and repetitive regions in order to finish the genomes. Assembly of the very large and internally repetitive gene for the biofilm-associated proteins Bap and BLP1 was a recurring problem. As these strains continue to be used for genetic studies and their genomes continue to be used as references in phylogenomics studies including core gene determination, there is value in improving the quality of their genome sequences. To this end, we re-sequenced two such strains that belong to the two major globally distributed clones of *A. baumannii*, using a combination of highly-accurate short-read and gap-spanning long-read technologies. Annotation of the revised genome sequences eliminated hundreds of incorrect CDS feature annotations and corrected hundreds more. Given that these revisions affected hundreds of non-existent or incorrect CDS features currently cluttering GenBank protein databases, it can be envisaged that similar revision of other early bacterial genomes that were sequenced using error-prone technologies will affect thousands of CDS currently listed in GenBank and other databases. These corrections will impact the quality of predicted protein sequence data stored in public databases. The revised genomes will also improve the accuracy of future genetic and comparative genomic analyses incorporating these clinically important strains.

**Data summary:**

1. The corrected complete genome sequence of *A. baumannii* AB307-0294 has been deposited in GenBank; GenBank accession number CP001172.2 (chromosome url - https://www.ncbi.nlm.nih.gov/nuccore/CP001172.2).
2. The corrected complete genome sequence of ACICU has been deposited in GenBank under the GenBank accession numbers CP031380 (chromosome; url - https://www.ncbi.nlm.nih.gov/nuccore/CP031380), CP031381 (pACICU1; url - https://www.ncbi.nlm.nih.gov/nuccore/CP031381) and CP031382 (pACICU2; url - https://www.ncbi.nlm.nih.gov/nuccore/CP031382).

**The authors confirm all supporting data, code and protocols have been provided within the article or through supplementary data files.**

## 5. Introduction

*Acinetobacter baumannii* is a Gram-negative bacterium that has emerged as an important opportunistic pathogen and is a research priority because of its high levels of resistance to antibiotics (1–3), desiccation, and heavy metals (4, 5). On a global scale, members of two clinically important clones, known as global clone 1 (GC1) and global clone 2 (GC2), have been responsible for the majority of outbreaks caused by multiply antibiotic resistant *A. baumannii* strains (1–3, 6–8). Whole genome sequencing (WGS) technologies have revolutionised the study of bacterial pathogens allowing the entire gene repertoire of bacterial strains to be determined and hence enabling the study of the relationships between outbreak strains with an unprecedented high resolution (9). However, accuracy is important.

The first 10 complete genomes of *A. baumannii* strains were reported between 2006-2012 (Table 1) and are still used as baseline in many studies of this microorganism (10–12). Except for three strains (AYE, TCDC-AB0715 and TYTH-1), all of the early *A. baumannii* complete genomes were sequenced using the 454-pyrosequencing technology and assembled using PCR. Pyrosequencing is known to generate frequent systematic sequencing errors, especially errors in the length of homopolymeric runs (13); and these errors lead to erroneous protein sequence (CDS) prediction, often associated with fragmentation of genuine open reading frames.

**Table 1.**
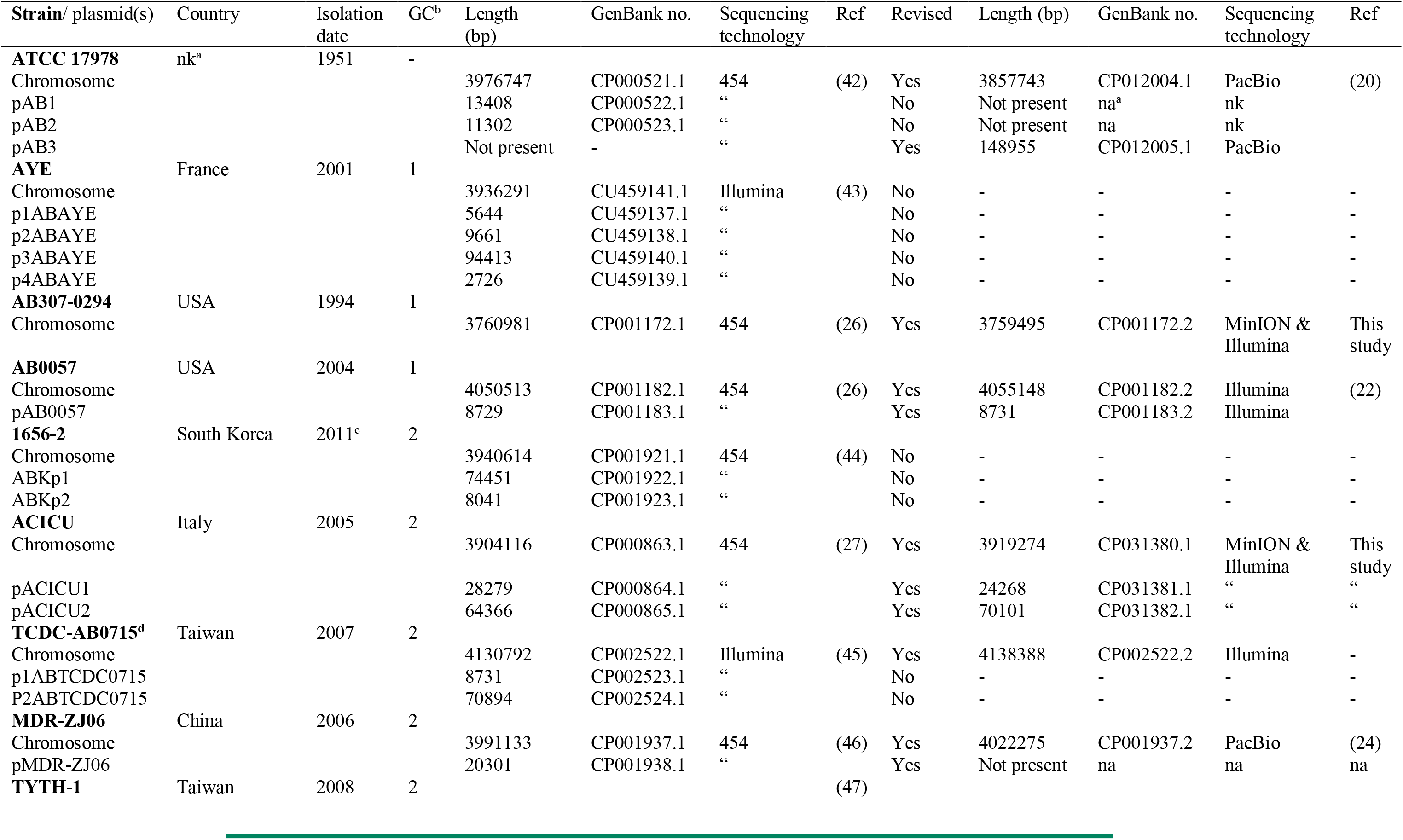
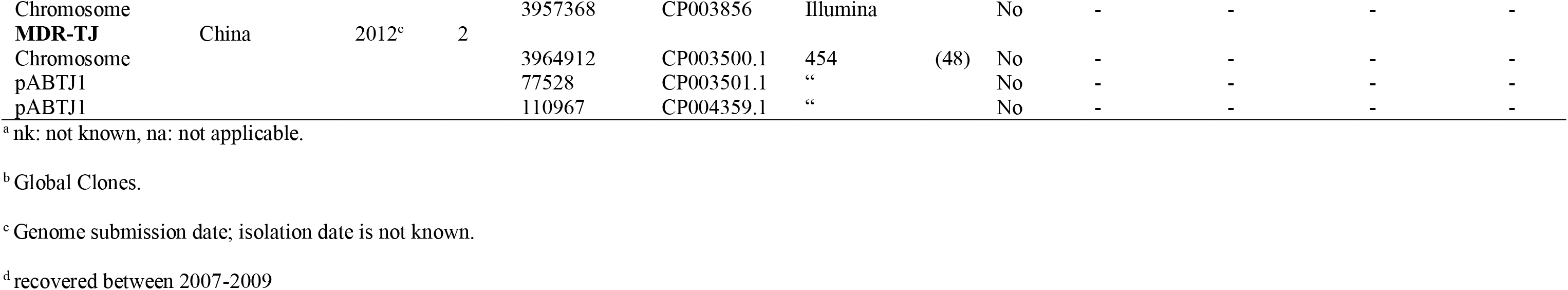
Properties of early *A. baumannii* completed genomes.

An additional problem in *A. baumannii* genomes determined using short read sequence data followed by PCR gap closure arises from the many short internal repeats present in the very large *bap* gene (~8-25 kbp), which is hard to assemble accurately. This gene encodes the biofilm associated protein Bap (14–17). The *bap* gene was originally cloned from AB307-0294 (GC1), and found to be 25,863 bp with a complex configuration of internal repeats (15). However, the size of the *bap* gene from a GC2 isolate was estimated at approximately 16 kbp (16). In another study, the length of Bap proteins predicted from *A. baumannii* genomes available in GenBank appeared to be highly variable, mainly due to different numbers of copies of the various repeated segments and the reading frame was often fragmented (17). The *blp1* gene, which is 9-10 kbp encodes a further very large protein that also has internal repeats and is associated with biofilm formation (17).

Newer sequencing technologies such as PacBio (Pacific Biosciences) and MinION (Oxford Nanopore Technologies, ONT) can generate much longer sequencing reads (9) allowing gaps to be spanned. MinION only assemblies are also prone to errors (18) but can be combined with high-accuracy Illumina short read data to produce very high quality finished genome assemblies (19). Long read sequence data has enabled a re-assessment of early completed *A. baumannii* genomes, including several of the first 10 to be sequenced (Table 1). For example, in 2016, ATCC 17978 was re-sequenced using PacBio. This revealed the presence of a 148 kb conjugative plasmid, pAB3, fragments of which were erroneously merged into the chromosome in the original 454-based assembly (20). This plasmid sequence brought together the parts of GI*sul2*, fragmented pieces of which had been randomly distributed in the chromosome in the original sequence (21). In 2017, we revised the 454-based genome sequence of the GC1 strain AB0057 using Illumina HiSeq technology, and found that hundreds of single base additions or deletions changed >200 protein coding features (CDSs) (22). An additional copy of the *oxa23* carbapenem resistance gene, located in Tn*2006*, was also found in the revised sequence of the chromosome (GenBank no. CP001182.2) (22, 23).

A recent revision of the 454-based genome of the GC2 strain MDR-ZJ06 using PacBio sequencing led to the correction of hundreds of CDS features and allowed reassessment of the localisation of important antimicrobial resistance regions (24). The position of transposon Tn*2009*, which carries the *oxa23* gene, was revised; and a region originally reported as a plasmid, but that had been predicted to be a chromosomally-located AbGRI3 type resistance island (25), was incorporated into the chromosome (CP001937.2) (24). In the revised genome, the two arrays of gene cassettes carrying antibiotic resistance genes in class 1 integrons are now in the correct resistance islands. These revisions exemplify the challenges encountered when relying solely on short read data to assemble bacterial genomes and highlight the extent and impact of pyrosequencing errors particularly on CDS predictions.

Two further *A. baumannii* strains for which only early 454-based genome sequences are available are the largely antibiotic susceptible isolate AB307-0294, recovered from the blood of a patient hospitalized in Buffalo, NY, in 1994 (26), and the extensively antibiotic resistant isolate ACICU recovered in 2005 from cerebrospinal fluid of patient in San Giovanni Addolorata Hospital in Rome, Italy (GenBank no. CP000863) (27). AB307-0294 was one of the first global clone 1 (GC1) strains to be completely sequenced (26) and has been extensively used in genetic studies (28–32). It belongs to CC1 (ST1) in the Institut Pasteur multi-locus sequence typing (MLST) scheme and to ST231 in the Oxford MLST scheme and carries the KL1 capsule genes and OCL1 at the outer core locus (33) (Table 1). Compared to other GC1 strains characterised to date, AB307-0294 is relatively susceptible to antibiotics (26), exhibiting resistance only to chloramphenicol (intrinsic) and nalidixic acid (acquired). It contains no plasmids.

ACICU was the first global clone 2 (GC2) isolate to be sequenced (27). It belongs to ST2 in the Institut Pasteur MLST scheme and carries the KL2 capsule genes and OCL1 at the outer core locus (34). ACICU is carbapenem resistant and also resistant to multiple antibiotics including third generation cephalosporins, sulfonamides, tetracycline, amikacin, kanamycin, netilmicin and ciprofloxacin (27). It contains two plasmids (27). However, we previously showed that the largest plasmid, pACICU-2, which was reported to include no resistance genes, is larger and contains the amikacin resistance gene *aphA6* in transposon Tn*aphA6*. The central segment of Tn*aphA6*, including the *aphA6* gene and one of the ISAba125 copies as well as a 4.7 kb backbone segment were missing in the original 454-based whole genome sequence (35).

Here, we report revised complete genome sequences for *A. baumannii* strains AB307-0294 (GC1) and ACICU (GC2), generated using MiSeq (Illumina) and MinION (ONT) sequence data. The new genome sequences correct hundreds of protein coding features generated by the presence of SNDs (single nucleotide differences) and small insertion/deletions of mainly 1-3 bases in the earlier 454 genome sequences.

## 6. Methods

### 6.1 Whole genome sequencing, assembly and annotation

Whole cell DNA was isolated and purified using the protocol described previously (1, 36). Libraries were prepared from whole cell DNA isolated from AB307-0294 and ACICU and were sequenced using Illumina MiSeq and ONT MinION. Paired-end reads of 150 bp and MinION reads of up to 20 kb were used to assemble each genome using the Unicycler software (v0.4.0) (19) using default parameters.

Protein coding, rRNA and tRNA genes were annotated using the automatic annotation program Prokka v1.13 (37). Regions containing antibiotic resistance genes and the polysaccharide biosynthesis loci, biofilm-associated proteins and genes used in the MLST schemes were annotated manually.

To compare previous CDS (≥ 25 aa CDS features) annotations with our new results, we wrote a script (github.com/rrwick/Compare-annotations) to quantify the differences. This script classifies coding sequences in the annotations as either exact matches, inexact matches, only present in the first annotation or only present in the second annotation. We also used the Ideel pipeline of Dr Mick Watson (github.com/mw55309/ideel) to assess the completeness of CDS annotated in each genome, by comparing the length of each CDS to that of its longest BLAST hit in the Uniprot database (as described in http://www.opiniomics.org/a-simple-test-for-uncorrected-insertions-and-deletions-indels-in-bacterial-genomes/).

## 7. Results and discussion

### 7.1 Revised genome of ACICU

ACICU, the first GC2 strain to be completely sequenced, contains AbaR2 in the chromosomal *comM* gene (27). As this AbaR resistance island type is more usually found in this location in GC1 strains (38) with an AbGRI1 type island in GC2 isolates (39), ACICU may represent a rare GC2 lineage. Here, the ACICU genome was re-sequenced using a combination of Illumina (MiSeq, 58x depth) and ONT (MinION, 253x depth) data. The new contiguous ACICU chromosomal sequence comprised 3,919,274 bp (GenBank no. CP001172.2), compared to 3,904,116 bp in the original submission (GenBank no. CP000863), making the revised chromosome 15,158 bp longer (Table 1). Most of the additional length in the revised chromosome was found to be due to a 11.2 kbp longer *bap* gene, which is just over 11 kbp and in 9 smaller orfs in the original sequence (locus_ids ACICU_02938 to ACICU_2946) as noted previously (17). In the revised genome sequence the *bap* gene is 22.2 kbp (BAP; locus_id DMO12_08904), mainly due to a large number of short strings of repeated sequences missing previously. Hence, some of the variation in length of *bap* reported previously (17) may be due to sequencing and assembly issues rather than genuine length variation in the *A. baumannii* population. The *blp1* gene in the original sequence (locus_id ACICU_02910) is 9510 bp and 9813 bp (locus_id DMO12_08811) in the revised genome.

The revised chromosome of ACICU differs from the original at 281 positions including 40 SNDs and 241 insertions or deletions of 1-3 bases (mostly in homopolymeric runs of As or Ts). The original annotation included 3677 protein-coding features (CDS features are ≥ 25 aa) whereas the revised genome annotation contains 3605 CDS features. Comparison of the CDS features indicated that only 3129 CDSs are identical between the two versions. The differences are mostly due to correction of open reading frames that were interrupted or fused due to errors in the 454 sequence and include 80 CDSs unique to the revised version and 142 CDS features in the original sequence that could not be found in the corrected chromosome. A further 396 CDS that are present in both versions are altered: of these, 8 have the same length, 285 are longer in the revised chromosome and 103 are shorter. Overall, 98.8% of all genes (n=3568) in the new assembly are within 5% of the maximum length of homologous proteins in Uniprot (i.e. the expected length), calculated using the ideel pipeline (see Methods). In the old assembly, only 95.8% (n=3494) of all genes are within 5% of this expected length. The distribution of length ratios is shown in Fig. 1A, highlighting a substantial population of CDS annotated in the old assembly that have lengths well below those of homologous proteins in Uniprot.

**Figure 1.**
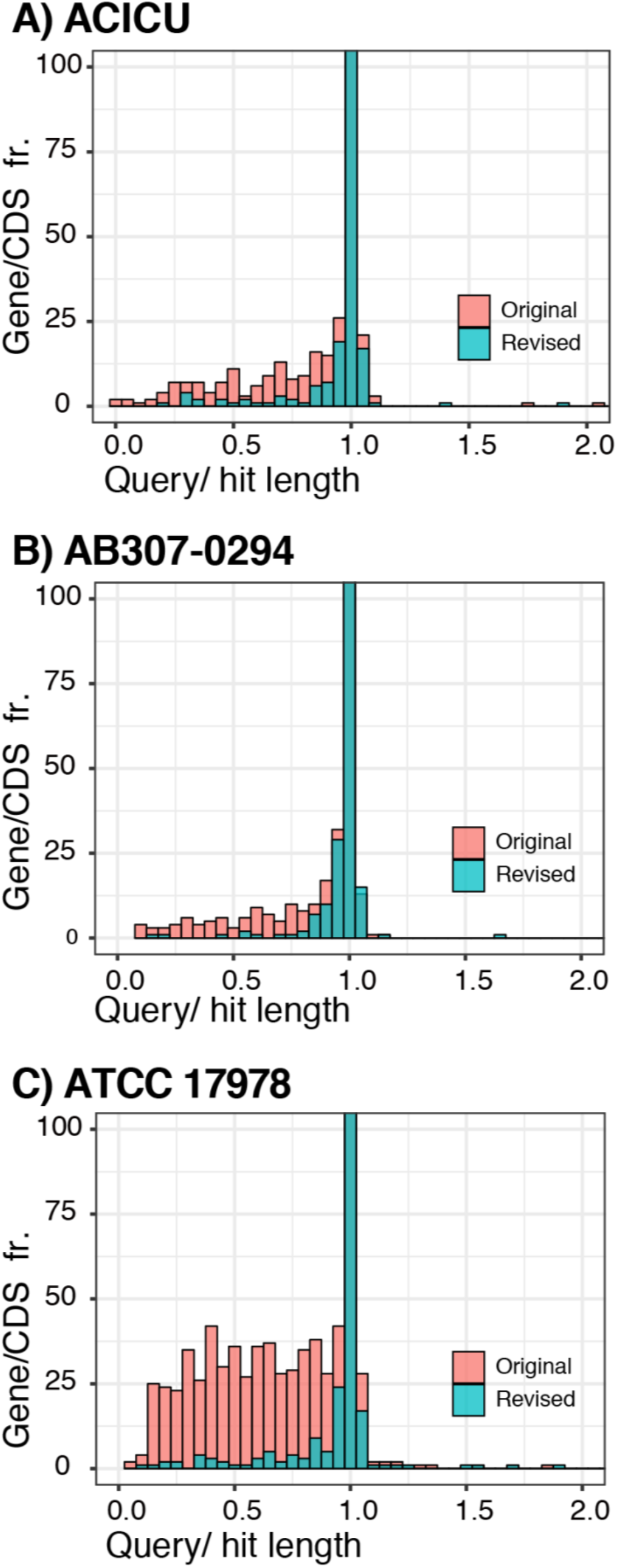
Histograms of CDS lengths relative to the length of the top hit in Uniprot, in the original vs revised genomes. A) ACICU GenBank accession no. CP000863.1 (original) and CP031380 (revised), B) AB307-0294 GenBank accession no. CP001172.1 (original) and CP001172.2 (revised), and C) ATCC 17978 GenBank accession no. CP000521.1 (original) and CP012004.1 (revised). The x-axis shows the ratio of coding sequence length to the length of the closest hit in the UniProt TrEMBL database. The y-axis shows gene frequency and is truncated at 100 (the centre bar extends to ~3000 genes). A tight distribution around 1.0 indicates that the assembly’s coding sequences match known proteins, supporting few indel errors in the assembly. A left-skewed distribution is characteristic of an assembly with indel errors which lead to premature stop codons.

ACICU carries two plasmids (Table 1), pACICU1 and pACICU2 (27), which encode the RepAci1 and RepAci6 replication initiation proteins (40). The original pACICU1 sequence (GenBank no. CP000864) is 28279 bp long and contains two copies of the carbapenem resistance gene *oxa58* while the revised pACICU1 (GenBank no. CP031381) is 24268 bp long and includes only a single *oxa58* copy. It lacks the region between the two IS*26* and one copy of IS*26* in the original sequence. The IS*26* mediated duplication may have been generated during growth in selective media. The original and revised pACICU1 sequences also differed by 3 SNDs, 6 single bp insertions and 1 single bp and 2 of 2 bp deletions. We previously used a PCR mapping strategy (35) to show that the *aphA6* gene and an additional ISAba125 as well as a 4.7 kb long backbone segment, located between two copies of a ~420 bp repeated segment, are missing from the original sequence of pACICU2, the larger plasmid of ACICU (35). Here, the long-read sequences generated for pACICU2 (GenBank no. CP031382) confirmed this. The revised plasmid sequence differs by 6 SND from pAb-G7-2 (GenBank no. KF669606.1), a conjugative plasmid from a GC1 isolated in Australia in 2003 reported previously (41).

### 7.2 Revised genome of AB307-0294

The AB307-0294 genome was also sequenced using a combination of Illumina (MiSeq, 63x depth) and ONT (MinION, 120x depth) technologies. The hybrid assembly resulted in a single 3,759,495 bp chromosome (GenBank no. CP001172.2) compared with 3,760,981 bp in the original genome (GenBank no. CP001172.1), making the revised genome 1486 bp shorter (Table 1). As with AB0057, the majority of differences were found to be additions or deletions of 1-3 bases, usually in “A” or “T” in homopolymeric runs of these nucleotides. The original annotation included 3427 CDS while the revised annotation contains 3458 (≥ 25 aa), of which 2937 CDSs are identical in the two versions. Corrections of insertion/deletion errors changed 354 reading frames leading to merging and splitting of CDS regions. Amongst these 354 CDS features, 286 CDSs in the revised genome are longer and 65 are shorter than the corresponding CDSs in the original annotation and 3 have the same length but differ internally. The revised genome also includes 136 novel CDS features, compared to the original sequence, while there are also 167 CDS in the old sequence that no longer exist in the revised genome again indicating the high impact of the errors caused by the use of 454-pyrosequencing technology. Overall, 98.9% of all genes (n=3387) in the new assembly are within 5% of the expected length, calculated using the ideel pipeline, versus just 96.4% (n=3336) in the old assembly (Fig. 1B).

The *bap* gene was 25863 bp (locus_id ABBFA_00771), the same length as reported originally (15) but 1067 bp shorter than the 26930 bp *bap* gene in the original genome sequence where it is split into two open reading frames (locus_id ABBFA_000776) and (locus_id ABBFA_000777). The revised genome was found to contain a 10089 bp *blp1* gene (ABBFA_00802), only 18 bp longer than that in the original sequence. Interestingly, both the original and revised genomes appear to be devoid of any insertion sequences (IS).

### 7.3 Revised genomes affect many predicted protein sequences

To date, 6 early *A. baumannii* genome sequences, including AB307-0294 and ACICU reported here, have been corrected and in each case the revised genome has resulted in correction of ~ 600 CDS features on average (20, 22). In each comparison of revised and original genome sequences, 100-150 new CDS features appeared, 150-200 CDSs disappeared and 150-200 CDSs changed. As the extent of errors had not been reported previously (20), we also compared the original (GenBank no. CP000521.1) and revised (GenBank no. CP012004.1) genomes of *A. baumannii* ATCC 17978. This revealed that the revised sequence has extensively re-ordered parts of the chromosome correcting a large number of inversions, insertion/deletions and other mis-assemblies. A striking difference between the two genomes is the inclusion in the original chromosome assembly of several large segments that in fact make up a 148 kb plasmid (pAB3) carrying the *sul2* sulfonamide resistance gene (GenBank no. CP012005). The misassembly issues precluded a simple alignment of the two chromosome sequences, but alignment of 14 separate chromosomal segments totalling 3843892 bp, revealed 334 SNPs as well as 635 deletions and 754 insertions of 1-3 bases, mainly “A”s or “T”s in runs of “A”s or “T”s. Overall, 3503 genes (98.2% of all genes) in the new assembly are within 5% of the expected length, calculated using the ideel pipeline, versus 3381 (86.4%) in the old assembly (see Fig. 1C). Hence, the original assembly was substantially flawed and should not be used in future. However, although the original study reported that ATCC 17978 contains two cryptic plasmids of 13 kb, pAB1 (GenBank no. CP000522.1) and 11 kb, pAB2 (GenBank no. CP000523.1) (42), the revised genome does not include either of these plasmids. This may be due to an assembly parameter setting to filter out the small contigs, which would remove pAB1 and pAB2, from the final assembly.

Granted the large effects observed on the length of *bap* and *blp* in ACICU using long read data, their sizes in original and revised genomes in the remainder of the first set of 10 (Table 1) were compared and significant differences were observed only where long read data was used in the revision. In the GC2 strain MDR-ZJ06 (GenBank accession no. CP001937), *blp1* (locus tag ABZJ_03096) is 9,812 bp in the revised genome (CP001937.2) versus 9,134 bp in the original sequence (locus tag ABZJ_03096). Further, *bap*, which is 7946 bp in the revised genome (locus_id ABZJ_03955) was split into 3 orfs, ranging in size from 2 to 2.5 kb, in the original sequence. In ATCC 17978, the *blp1* gene is not present in either the original or the revised genome. However, the *bap* gene, which was split into two open reading frames (locus_id A1S_2696; 6306 bp and A1S_2724; 1161 bp) and separated by 41 kbp in the original sequence is now in a single orf (locus_id ACX60_04030; 6225 bp) in the revised genome and 842 bp shorter compared to those in the original genome.

## 8. Conclusions

The revised genome sequences of AB307-0294 and ACICU will underpin more accurate studies of the genetics and genomic evolution of related *A. baumannii* strains belonging to GC1 and GC2.

This work highlights the need to review and revise early bacterial genomes sequenced using short read data and assembled with (or sometimes without) PCR to join contigs. Special attention needs to focus on the genomes determined using the 454-pyrosequencing technology in order to correct predicted protein sequences.

Long read data, such as those generated by PacBio and ONT (MinION) technologies, allows for complete genome assembly without manual intervention. While assembling long read data alone can result in sequence errors and failure to detect small plasmids, hybrid assembly (using both short and long reads) can produce assemblies that are both complete and accurate. However, repetitive sequences in the genome, such as the genes encoding Bap and BLP1, are difficult to perfect even with hybrid assembly, so variations in these regions should be interpreted with caution.

Finally, as the original GenBank entries are replaced by revised genomes, there is a need to eliminate non-existent and incorrect predicted protein sequences in order to simplify the already complex task of protein sequence searches. It can be assumed that this problem is not only limited to *A. baumannii* genomes as many bacterial species so far have been sequenced using the 454-pyrosequencing technology.

## 9. Author statements

### 9.1 Authors and contributors

Conceptualization, RMH, MH; Data curation, MH, RW; Formal analysis, MH, RW, KEH, RMH; Funding, RMH, KEH, MH; Investigation, MH, RW, LJ; Resources, KEH; Visualization, MH, RW, KEH; Manuscript preparation, original draft, MH and RMH; review and editing RMH, MH, RW, KEH.

### 9.2 Conflicts of interest

The authors declare that there are no conflicts of interest.

### 9.3 Funding information

This work was supported by NHMRC grant GNT1079616 to RMH. MH is supported by the Chancellor’s Postdoctoral Research Fellowship (CPDRF PRO17-4005) from the University of Technology Sydney, Australia. KEH is supported by a Senior Medical Research Fellowship from the Viertel Foundation of Australia.

### 9.4 Consent for publication

Not applicable.

### 9.5 Ethical Approval

No human or animal experimentation is reported.

## 9.6 Acknowledgements

We would like to thank Prof. Thomas A. Russo, State University of New York, Buffalo, New York, USA, for kindly providing AB307-0294 and Prof. Alessandra Carratoli, Istituto Superiore di Sanità, Rome, Italy, for supplying ACICU.

